# Patient stratification of clear cell renal cell carcinoma using the global transcription factor activity landscape derived from RNA-seq data

**DOI:** 10.1101/829796

**Authors:** Yanyan Zhu, Shundong Cang, Bowang Chen, Yue Gu, Miaomiao Jiang, Junya Yan, Fengmin Shao, Xiaoyun Huang

## Abstract

Clear cell renal cell carcinoma represents the most common type of kidney cancer. Precision medicine approach to ccRCC requires an accurate stratification of patients that can predict prognosis and guide therapeutic decision. Transcription factors are implicated in the initiation and progression of human carcinogenesis. However, no comprehensive analysis of transcription factor activity has been proposed so far to realize patient stratification. Here we propose a novel approach to determine the subtypes of ccRCC patients based on global transcription factor activity landscape. Using the TCGA cohort dataset, we identified different subtypes that have distinct upregulated biomarkers and altered biological pathways. More important, this subtype information can be used to predict the overall survival of ccRCC patients. Our results suggest that transcription factor activity can be harnessed to perform patient stratification.

## Introduction

Kidney cancer is one major solid cancer, with an estimated 403262 new cases and 175098 deaths in 2018[1]. Renal cell carcinoma accounts for around 90% of kidney cancer, with clear cell renal cell carcinoma being one of the most frequent subtypes. Despite recent advances in targeted therapy and immunotherapy, there is still a large gap to be filled for the clinical management of patients diagnosed with clear cell renal cell carcinoma.

Transcription factors are implicated in the initiation and progression of human carcinogenesis. The c-myc oncogene encodes a transcription factor that regulates the transcription of hundreds of genes. It has been well established that transcription factors are situated in the hubs of a complex network shaping the hallmarks of human cancer. Thus, it is intriguing to develop strategies to target transcription factors, especially a combination targeting strategy designed to destroy cancer cell dependency[2]. Small molecules with the capacity to reactivate mutant p53 are now being evaluated in clinical trials[3]. Furthermore, transcription factors can be harnessed to develop combination strategy aiming to defeat the intratumor heterogeneity. For example, combination treatment with STAT3 and BCL6 inhibitors were shown to reduce the growth of xenografted tumors[4].

An accurate determination of the subtypes of clear cell renal cell carcinoma is essential to develop personalized therapy for ccRCC patients. ccRCC has been stratified previously with genomic and transcriptomic information[5, 6]. However, no studies have focused on transcription factor. Considering the fact that various transcription factors play an important role in metabolic rewiring[7], growth and metastasis[8] of ccRCC cells, it is tempting to approach the molecular stratification of ccRCC using the landscape of transcription factor activity. Here we construct a comprehensive atlas of transcription factor profiles using dataset available in the TCGA cohort comprising of 603 samples.

## Method

### Download TCGA level 3 data

TCGA level 3 expression data were downloaded from the Pan-Cancer Atlas datasets hosted at Genomic Data Commons (https://gdc.cancer.gov/node/905/). In total, gene expression profiles for 531 ccRCC primary cancer tissues and 72 solid normal tissues were extracted from the Pan-Cancer Atlas.

### Transcription factor scoring

For each transcription factor, the target genes with known regulation modes were extracted from TTRUST database[9], resulting in a list of genes activated by the transcription factor and a list of genes repressed by the transcription factor. The ratio between the median expression level of activated target gene and the median expression level of repressed target gene was calculated for each transcription factor and log2 transformed to obtain a final transcription factor score.

### k-means clustering

k-means clustering was performed to unbiasedly analyse the structures in the dataset. Only those transcription factors which have no NAs were included in the clustering analysis. In total, 238 transcription factors were analysed. Subpopulations of patients were estimated using only the transcription factor score matrix. The optimal number of k was determined by gap statistic.

### Differential expressed genes

The unique biomarkers characterizing one cluster was determined by statistical test comparing samples within that particular cluster and the rest of all other samples. “bonferroni” correction was employed to reduce false hits in multiple comparisons. Adjusted *p-*value less than 0.05 was considered as statistically significant. Genes with fold change greater than 1.5 were selected as candidate genes for downstream analysis.

### Gene list analysis

Gene list analysis was performed using metascape[10]. Four major steps were included in the pipeline: ID conversion, Annotation, Membership determination and Enrichment analysis. Differentially expressed gene list was input into the online tool and analysed using default parameters. Statistically enriched terms were identified and filtered. Remaining significant terms were then hierarchically clustered into a tree based on Kappa-statistical similarities among their gene memberships. A subset of representative terms was selected from the full cluster and converted into a network layout. More specifically, each term is represented by a circle node, where its size is proportional to the number of input genes fall into that term, and its color represent its cluster identity. Terms with a similarity score > 0.3 are linked by an edge (the thickness of the edge represents the similarity score). The network is visualized with Cytoscape (v3.1.2) with “force-directed” layout and with edge bundled for clarity. One term from each cluster is selected to have its term description shown as label.

### Protein-protein interaction

All protein-protein interactions (PPI) among each input gene list were extracted from PPI databases and formed a PPI network. GO enrichment analysis was applied to the original PPI network and its MCODE network components to assign biological “meanings”, where top three best *p*-value terms were retained. All input gene lists were also merged into one list and resulted in a PPI network. MCODE components were identified from the merged network. Each MCODE network is assigned a unique color. Network nodes are also displayed as pies. Color code for pie sector represents a gene list.

### Survival analysis

Kaplan Meier analysis was used for survival curve analysis. It measures the percent of patients surviving with time.

### Statistical method

All statistical analyses were performed using R. p-value < 0.05 was considered statistically significant. All plots were generated with R.

## Result

### Establishment of an analytical framework to quantify the global transcription factor activity

One typical transcription factor regulates the expression of hundreds of downstream gene either positively or negatively. To quantify the activity of one transcription factor, the expression of both the activated genes and the repressed genes should be taken into consideration. A metric was devised to incorporate both into one score, through which a comprehensive landscape of transcription factor activity was established (**Figure 1A**). Globally, all transcription factors were grouped into “activated” and “repressed” states. To further gain insights into the patterns in the transcription factor landscape, the correlation matrix was visualized with a heatmap. It was shown that all samples fell into two apparent groups and the larger group harbored finer structures (**Figure 1B**).

**Figure 1.**
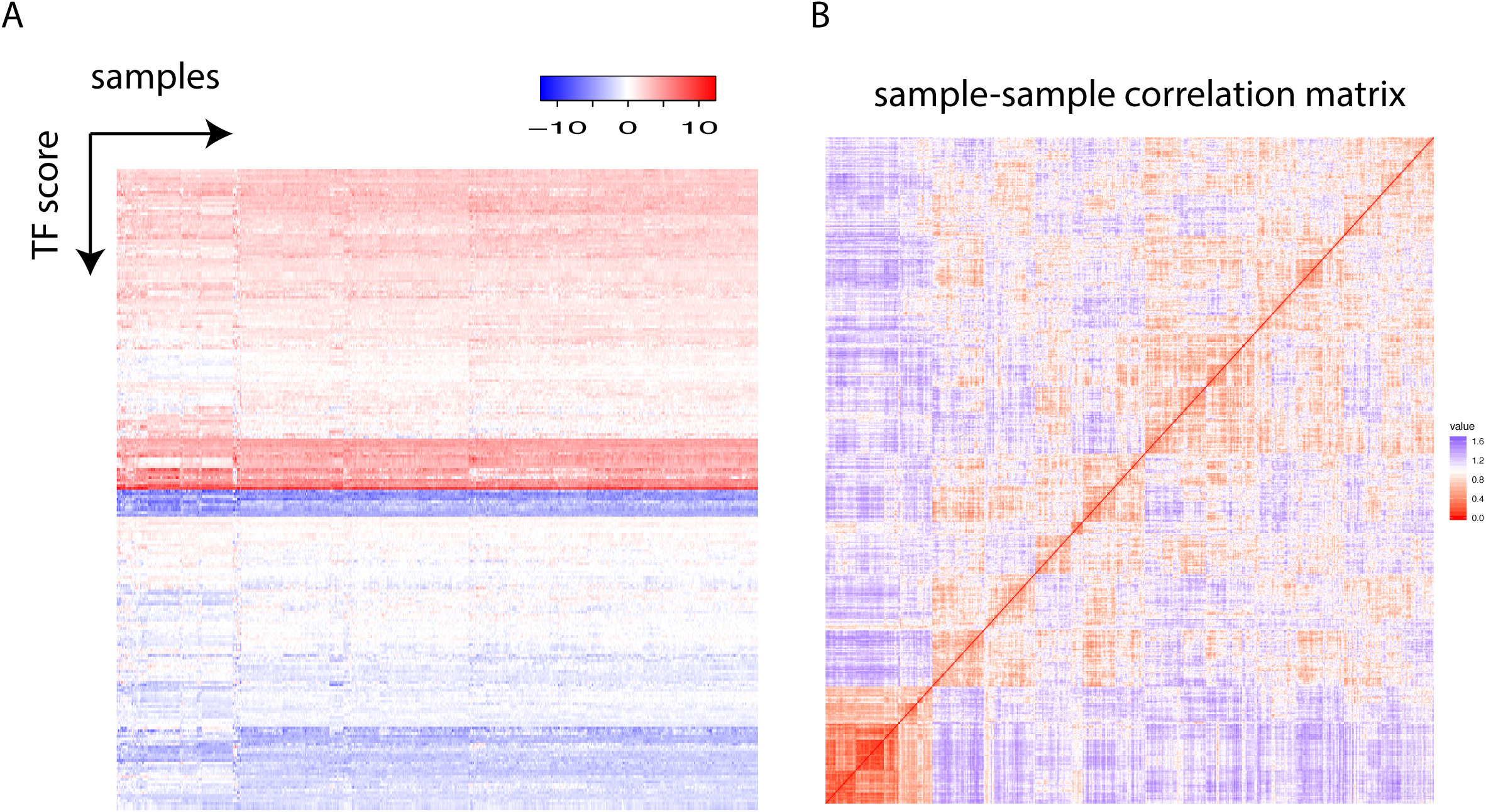
(A) Each row represents one transcription factor and each column stands for one individual. The transcription factor score is plotted. (B) The sample-sample correlation matrix is visualized as a heatmap.

### Patient stratification based on global transcription factor activity

To estimate the number of clusters in the samples, gap statistic was employed. It was found that four clusters could contain more than 88% of the information in the TCGA dataset (**Figure 2A**). To identify the clusters, k-means clustering was performed using all the samples in the TCGA dataset, which included 531 cancer samples and 72 normal tissues (**Figure 2B**). Of note, all normal samples were grouped together as Subtype_3. No normal samples were in the other clusters, while there were also some cancer samples in Subtype_3. Those cancer samples were normal like in the transcription factor activity.

**Figure 2.**
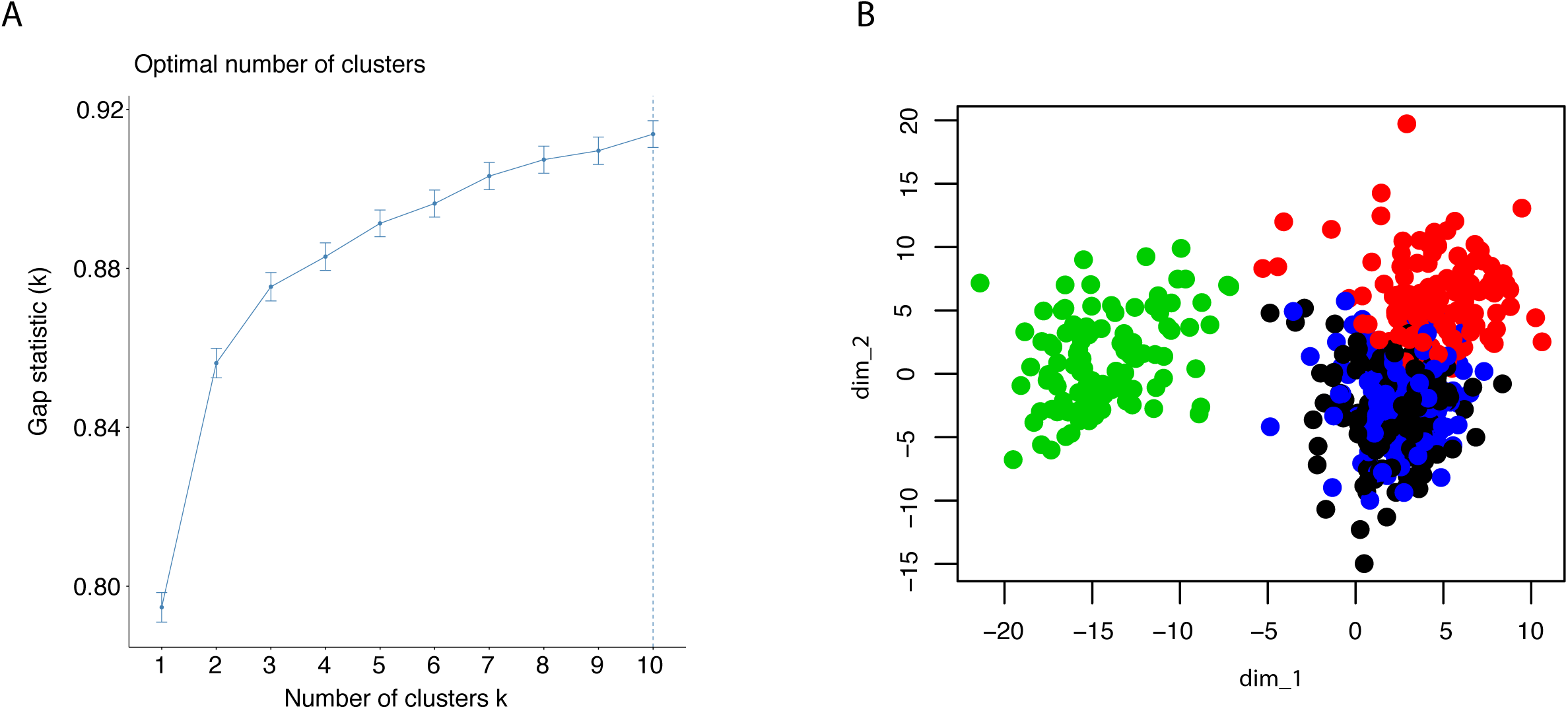
(A) Gap statistic was shown up to 10 clusters, to determine the optimal k for k means clustering. (B) The result of k means clustering was plotted for k = 4, using all individuals.

Next, differential gene expression analysis was performed using statistical test. Each subtype identified previously was compared with the rest of all subtypes, resulting a list of genes whose expression is characteristic for that subtype. Top 10 up-regulated genes for each subtype were visualized with heatmap (**Figure 3A**). The up-regulated genes with the Subtype_3 included genes essential for the normal functionality of kidney cells, while the other three subtypes over-expressed genes with known roles in carcinogenesis. Alteration of angiogenesis is common in ccRCC and this typically involves the VEGF (**Figure 3B**). Not surprising, cancer clusters expressed significantly higher level of VEGF. Besides an upregulated angiogenesis, expression of PD-1 on cancer cells were correlated with the ability of cancer cells to evade the immune systems. Consistently with previous observations, cancer clusters upregulated the expression of PD-1(**Figure 3B**).

**Figure 3.**
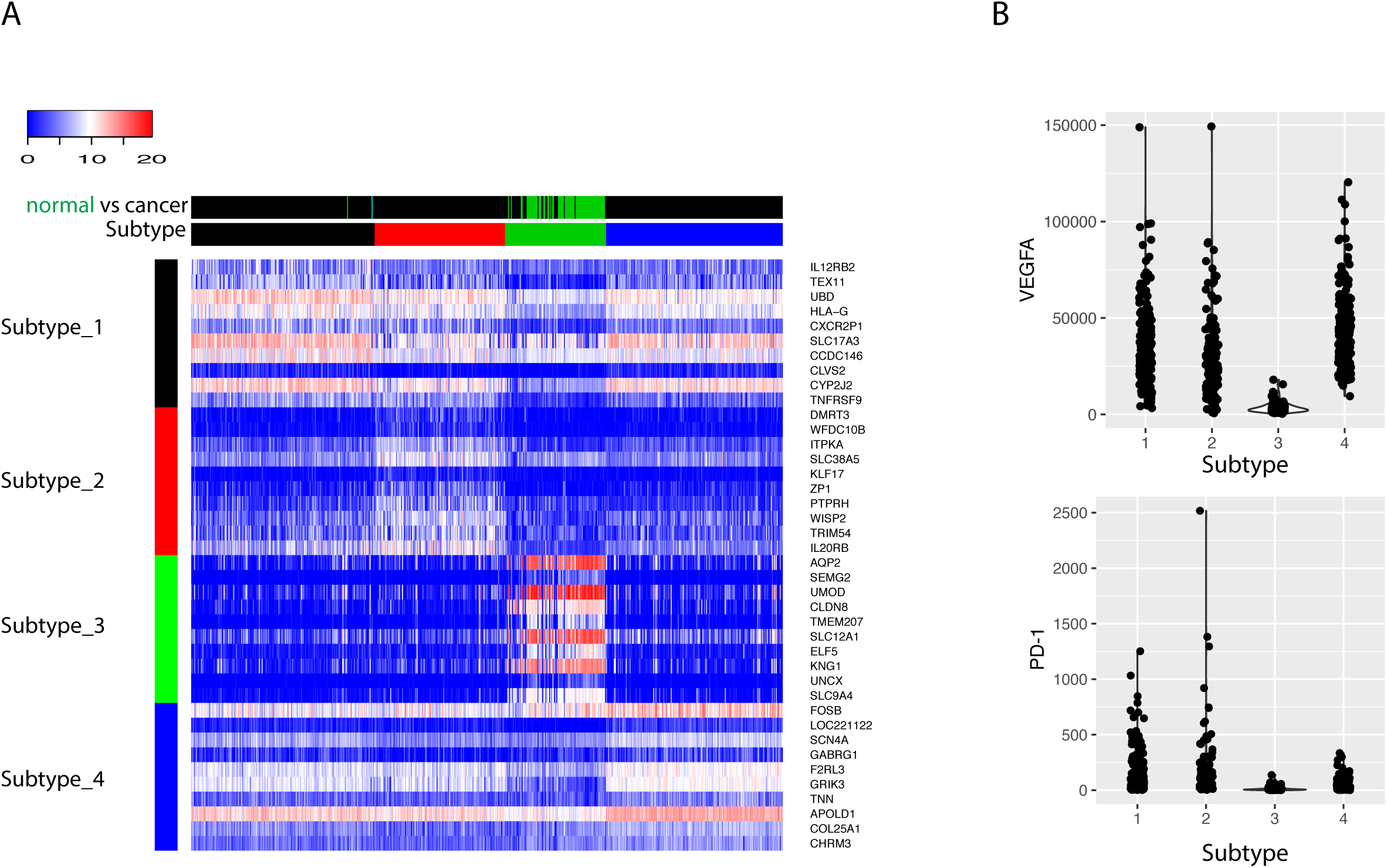
(A) The top 10 biomarkers upregulated in each subtype were shown as heatmap. Each row represents one gene, and each column stands for one sample. The column side color bar labels the information about the sample. In Subtype bar, Subtype_1, Subtype_2, Subtype_3 and Subtype_4 are colored as black, red, green and blue respectively. In normal vs cancer bar, cancer tissue is labeled as black and normal tissue is labeled as green. (B) The expression of VEGFA and PD-1 was shown for the four subtypes: Subtype_1, Subtype_2, Subtype_3 and Subtype_4.

### Different patient groups have distinct pathway alterations

To investigate what pathways or biological processes were uniquely altered for the different subtypes, GO analysis was performed. Interestingly, the Subtype_3 stood out while the other three subtypes were grouped together after hierarchical clustering of the enriched GO terms (**Figure 4A**). GO terms uniquely enriched in the Subtype_3 included lipid biosynthetic process, TCA cycle, monovalent inorganic cation homeostasis, mitochondrion organization, metabolism of vitamins and cofactors, suggesting the downregulation of the normal kidney functions in the other subtypes. The fact that only a few cancer samples were in Subtype_3 indicates that most ccRCC cancer cells tune down normal processes exerted by normal kidney cells during carcinogenesis. Subtype_1 and Subtype_2 have altered T cell signaling pathways, such as adaptive immune response, cytokine production, cell activation in the immune system and T cell selection. It was consistent with some previous reports showing that cancer rewired the tumor microenvironment to become immune suppressive environment. Subtype_4 preserved some of the functionality of normal kidney cells, such as regulation of ion transport. Of note, Subtype_2 have significantly altered cell division, mitotic nuclear division and cellular response to lipids, unlike any other subtypes. Globally, regulation of cell adhesion was altered in Subtype_1 and Subtype_2 (**Figure 4B**).

**Figure 4.**
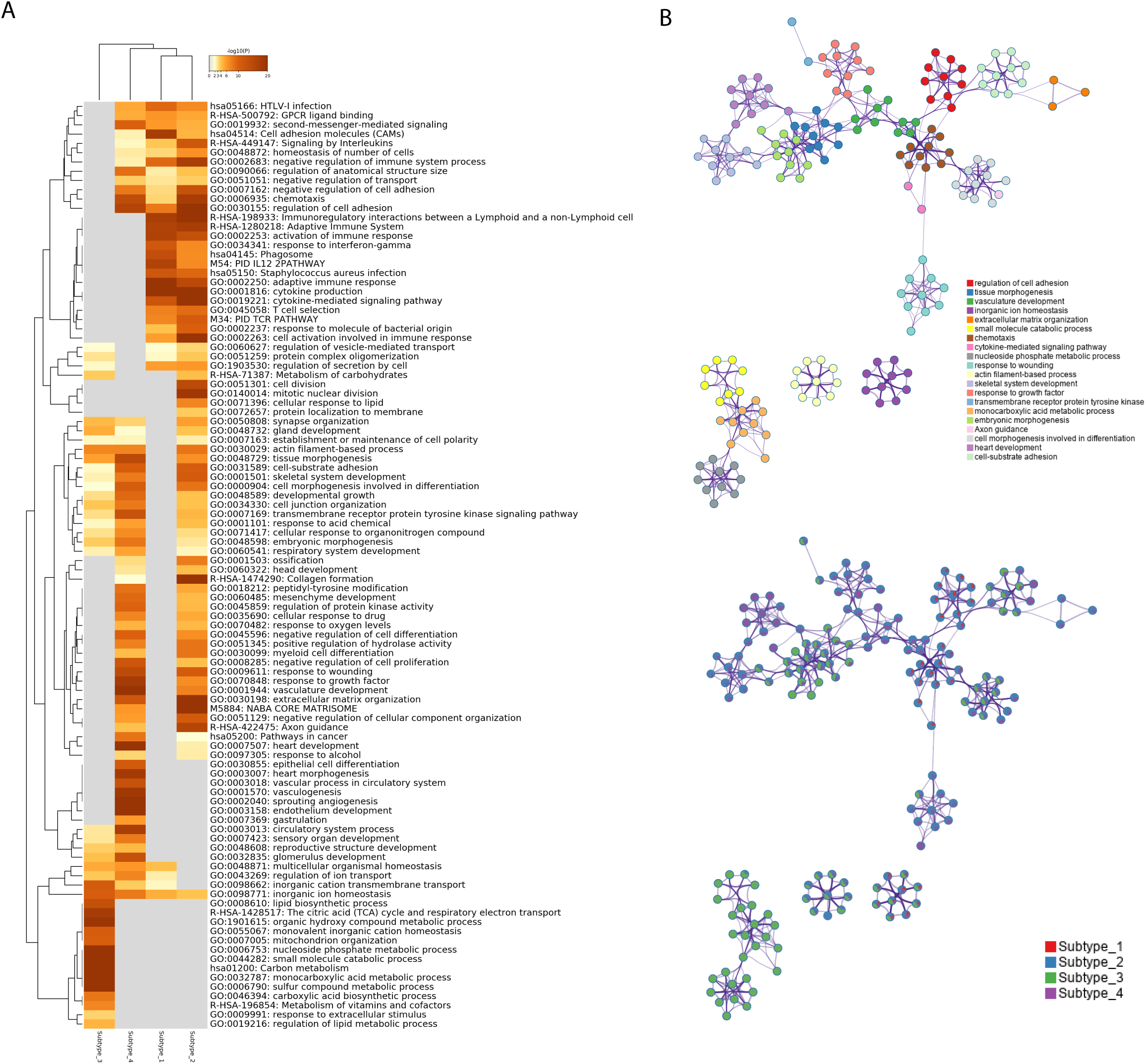
(A) Statistically enriched terms were identified and then hierarchically clustered into a tree based on Kappa-statistical similarities among their gene memberships. The heatmap cells are colored by their p-values, white cells indicate the lack of enrichment for that term in the corresponding gene list. (B) A subset of representative terms from the full cluster was selected and converted into a network layout. Each term is represented by a circle node, where its size is proportional to the number of genes satisfying that term and its color represents its cluster identity. Terms with a similarity score > 0.3 are linked by an edge where the thickness of the edge indicates the magnitude of similarity. The same enrichment network has its nodes displayed as pies. Each pie sector is proportional to the number of hits originated from a gene list.

### Immunity related networks are up-regulated in Subtype_1 and Subtype_2

Using public databases with protein-protein interaction information, MCODE motifs in the regulated genes were extracted (**Figure 5A**). Subtype_1 and Subtype_2 shared MCODE networks in T cell signaling pathways, suggesting an important role played by the immune systems. To consider the contribution of genes for the identified MCODE, it was found that Subtype_2 was the only cluster that contributes to all identified MCODE (**Figure 5B**). Besides two MCODE related to immunity, there was one MCODE consisting of three players: NOTCH1, NOTCH3, and SMARCD3. SMARCD3 is an essential component of the SWI/SNF chromatin-remodeling complex, which epigenetically modulates the expression of downstream genes.

**Figure 5.**
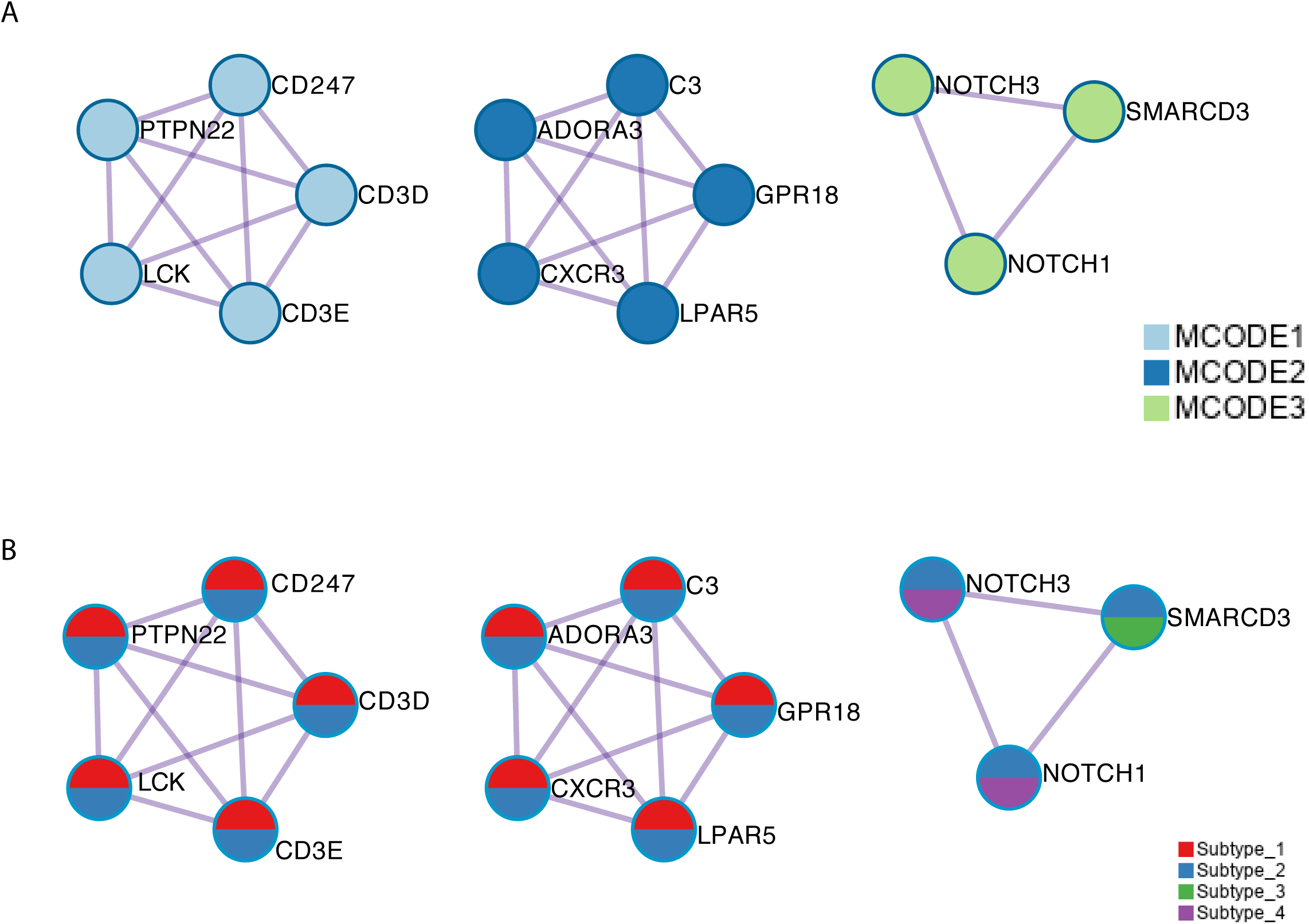
(A) The MCODE components were identified from the merged protein-protein interaction network. Each MCODE network is assigned a unique color. (B) The same MCODE networks were displayed as in (A), where network nodes shown as pies. Color within pie sector indicates the subtype origin.

### Prognostic value of the proposed transcription factor stratification

The transcription factor landscape is sufficient to divide the cancer patients into subtypes, with distinct biomarker and altered pathways. We then ask whether different subtypes have different overall survival. The survival information was downloaded to devise the survival curves for different subtypes(**Figure 6**). Here we used only the cancer samples in the dataset. For the three cancer only subtypes, Subtype_2 has the worse OS as compared with all the other subtypes. Subtype_4 has the longest OS in the three cancer only subtypes. Interestingly, the normal like Subtype_3 did not have significantly better OS compared with Subtype_1 and Subtype_4.

**Figure 6.**
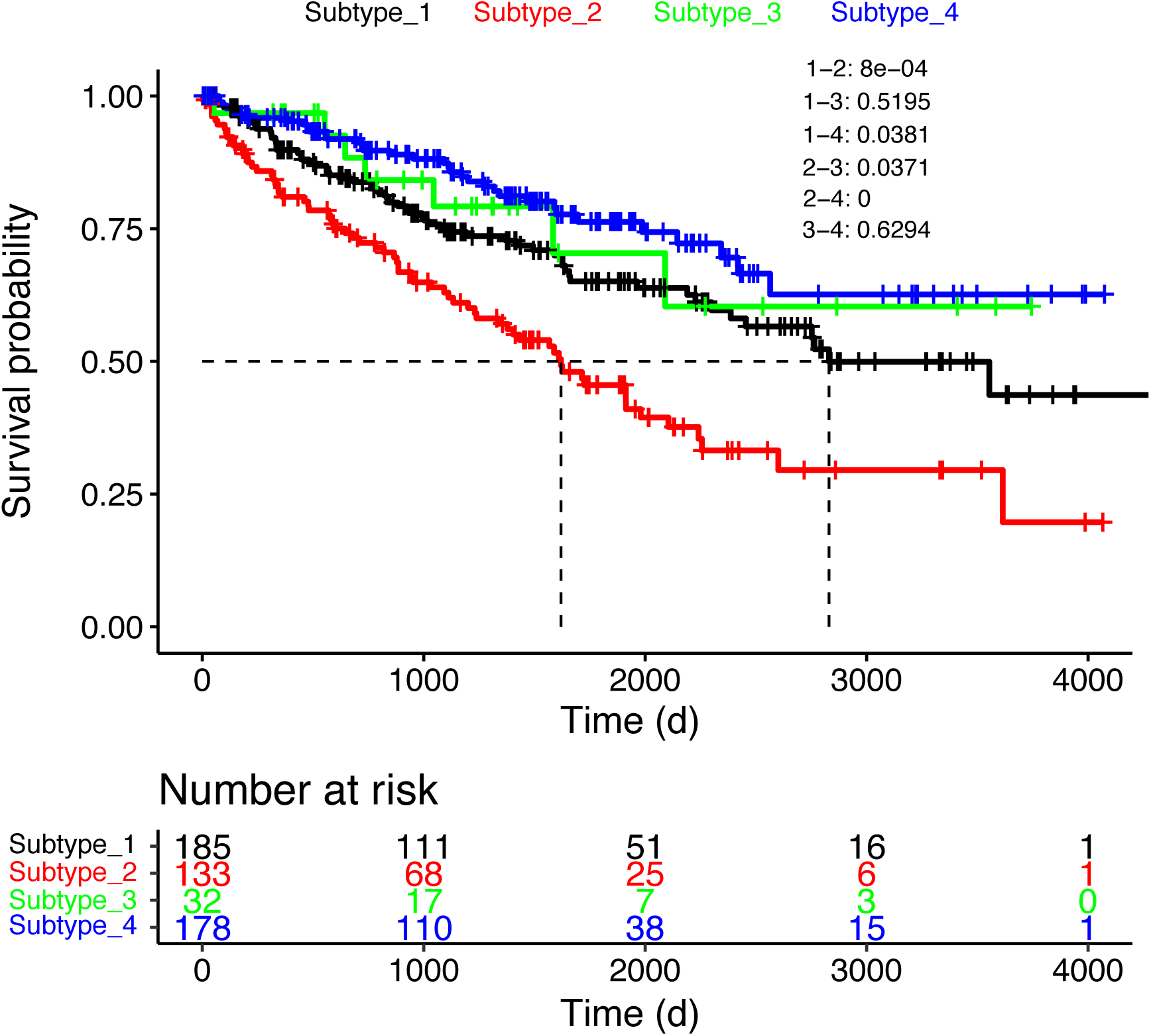
The survival curves were shown for the four subtypes. Only cancer samples were used for the analysis.

## Discussion

Patient stratification is key to personalized therapy of cancer patients. One milestone paper[11] divided ccRCC into two major subtypes: clear cell type A (ccA) and clear cell type B (ccB). Subtype ccB was correlated with poor survival compared with subtype ccA. Pathway analysis suggested that ccA group overexpressed genes associated with hypoxia, angiogenesis and metabolism, while ccB group overexpressed genes involved in EMT, cell cycle and wound healing. Unsupervised clustering of the TCGA ccRCC cohort revealed four stable subtypes using RNA expression data designated m1-m4[6]. The subtype m1 corresponded to the ccA group mentioned earlier while the subtype m2 and m3 corresponded to the ccB group. The remaining subtype m4 represented a previously uncharacterized subtype with a frequency of around 15%. The m1 subtype was characterized by upregulation of genes involved in chromatin remodeling.

Transcription factors regulates the transfer of genetic information from DNA to RNA. Traditionally, transcription factors were considered as undruggable. However, a shift of paradigm is realized as multiple approaches to modulate the activity of transcription factors have been demonstrated both preclinically and clinically[12]. Targeting transcription factors can be an effective strategy against RCC. It has been demonstrated that STAT3 inhibitor WP1066 inhibited the growth of renal cancer cell lines or xenografted renal cancer cells[13]. Checkpoint inhibition is emerging as a promising therapy for ccRCC patients, with a subset of patients responding to anti-PD-1 monotherapy extremely well. Gene expression analysis uncovered an altered transcriptional output in Janus kinase-signal transducers and activators of transcription (JAK-STAT) signaling[14]. It was shown that alteration of transcription factor activity can influence the response to checkpoint inhibitors.

Large cancer genome programs such as TCGA has generated an enormous amount of data that is publicly available to the cancer research community. Mining those datasets have enabled novel insights into cancer biology[15-19]. Our analysis suggests that predictive models can be derived from those massive data and used to assign patients to distinctive subtypes.

Our study can be extended by proof-of-concept experiments to establish the feasibility of predicting drug sensitivity based on the proposed patient stratification. For example, experiments can be carefully designed to investigate the effect of CDK4/CDK6 inhibitors in treating Subtype_2 cancer patients, using in vitro cancer cell line model, ex vivo or in vivo cancer models. One popular model to test is the patient derived organoids[20]. Subtyping can be performed with transcriptomic analysis using RNA-seq and the organoids can be perturbed with drug treatments to show proof-of-concept that optimal treatment strategy can be predicted with transcription factor activity based ccRCC subtyping.

## Conclusion

In conclusion, we have resisted the molecular stratification of clear cell renal cancer by integrative analysis of multiple publicly available datasets with a focus in the landscape of transcription factor activity. We demonstrated how a deep understanding of ccRCC subtypes can be obtained by a comprehensive and integrative reanalysis of publicly available datasets.

## Author Contribution

YZ, BC, MJ, and XH performed the analysis. YZ, SC, YG, JY and XH interpreted the results. XH and FS conceived and supervised the study. All authors read and approved the manuscript.

## Acknowledgement

This study is partially supported by Henan Provincial Medical Science and Technology Advancement Program (201702161).

